# Telomerase mRNA Reduces Radiation-induced DNA Damage of human skin

**DOI:** 10.1101/2025.02.01.636031

**Authors:** David F. Chang, Shuang Li, Karem Court Pinto, Thi Kim Cuc Nguyen, Vrutant V. Shah, Elisa Morales, Jack Carrier, Andrew T. Ludlow, Kristopher W. Brannan, Aldona J. Spiegel, Olmsted-Davis, Biana Godin, Anahita Mojiri, John P. Cooke

## Abstract

Over four million people undergo radiation therapy annually in the United States. Among these, more than 90% experience varying degrees of radiation-induced skin injury. Despite the enormity of the problem, there is currently no FDA-approved agent to prevent or treat skin damage caused by ionizing radiation. In the current study, ionizing radiation induced dose-dependent genomic and mitochondrial DNA damage, leading to apoptosis in primary cutaneous cells. Prior treatment with mRNA encoding telomerase reverse transcriptase (TERT) substantially reduced radiation-induced DNA damage in human primary skin cells and tissues. Mechanistically, TERT mRNA pretreatment enhances DNA repair, reduces mitochondrial ROS, and decreases apoptosis without extending telomere length during the experimental period, suggesting a non-canonical function of TERT to accelerate the cellular recovery from radiation. These findings highlight a potential therapeutic approach for preventing radiation-induced skin injury.

**Graphic abstract:** 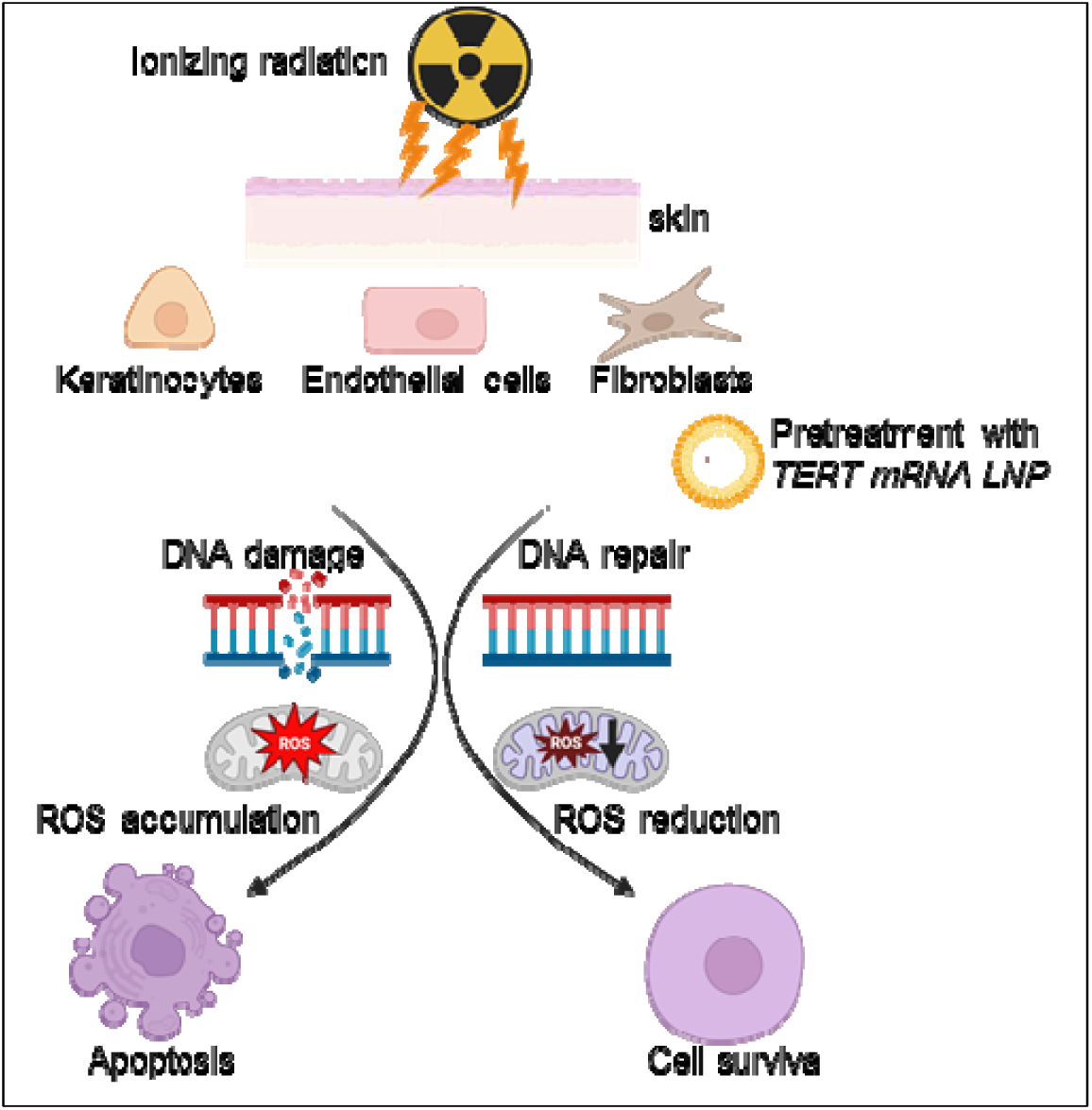

## Introduction

Radiotherapy induces DNA damage to destroy or halt the growth of malignant cells. Whereas it is effective against many types of cancer, the collateral damage to normal tissues poses a significant challenge ^1^. Notably, approximately 85–95% of patients’ undergoing radiotherapy experience varying degrees of cutaneous radiation injury, manifesting as erythema, dry and wet desquamation, secondary ulceration, and even cutaneous cancer. The cutaneous damage can severely compromise quality of life, particularly in those patients undergoing treatment for cancer involving the breast or head and neck. There is an unmet need for interventions that can mitigate these adverse effects while maintaining treatment efficacy.

Telomerase holoenzyme is a complex of several proteins including telomerase (TERT) which has reverse transcriptase activity. The canonical function of telomerase is to maintain telomere integrity, which is well described in both telomere extensions and maintenance, protecting cells from accumulation of DNA damage. Previously, we observed that, in human skin cells derived from adult patients, a single treatment with human telomerase (hTERT) mRNA could reverse DNA damage, increase proliferation, and enhance engraftment in a pre-clinical wound model ^2^. These observations were consistent with our prior work showing that in replicatively aged human cells, or in cells derived patients with genetically accelerated aging, treatment with TERT mRNA extends telomeres, normalizes cell and nuclear morphology, restores cellular functions and proliferation, normalizes the transcriptional profile, and reduces markers of senescence, oxidative stress, inflammatory cytokines, and DNA damage ^2,3–5^. Furthermore, in human endothelial cells derived from patients with genetically accelerated aging, hTERT mRNA increased the expression of genes that participate in repair of genomic DNA damage (such as FANCC, PARPBP, and RECQL5) ^5^. Although telomerase is well known to repair and to extend telomeres, it is less widely recognized that telomerase can also repair genomic DNA damage ^6^. Accordingly, the current study was driven by the hypothesis that TERT mRNA might protect human skin cells from DNA damage induced by clinically relevant levels of radiation.

In this study, we find that in primary human skin cells, as well as human skin explants, radiation induced DNA damage, mitochondrial dysfunction and apoptosis are attenuated by the expression of human TERT. Intriguingly, this appears to be due to a non-canonical function of hTERT, as the beneficial effects of hTERT mRNA delivery are seen in the absence of telomere extension indicating that telomerase holoenzymes are not required. Our observations may be useful in the development of a new therapeutic avenue for radiation-induced skin damage.

## Results

### Dose-dependent DNA damage response in cutaneous tissues following *ex vivo* ionizing radiation (IR) exposure

To evaluate whether ionizing radiation induces DNA damage in human cutaneous tissue, we exposed *ex-vivo* skin tissue to 2, 5 or 10 Gy of ionizing radiation and performed immunofluorescence staining for the DNA damage marker γH2AX, 2 hours post-exposure. We observed a dose-dependent increase of γH2AX signal without structural disruption of the skin (Fig 1A-E). The increase in γH2AX levels was most notable in keratinocytes.

**Figure 1.**
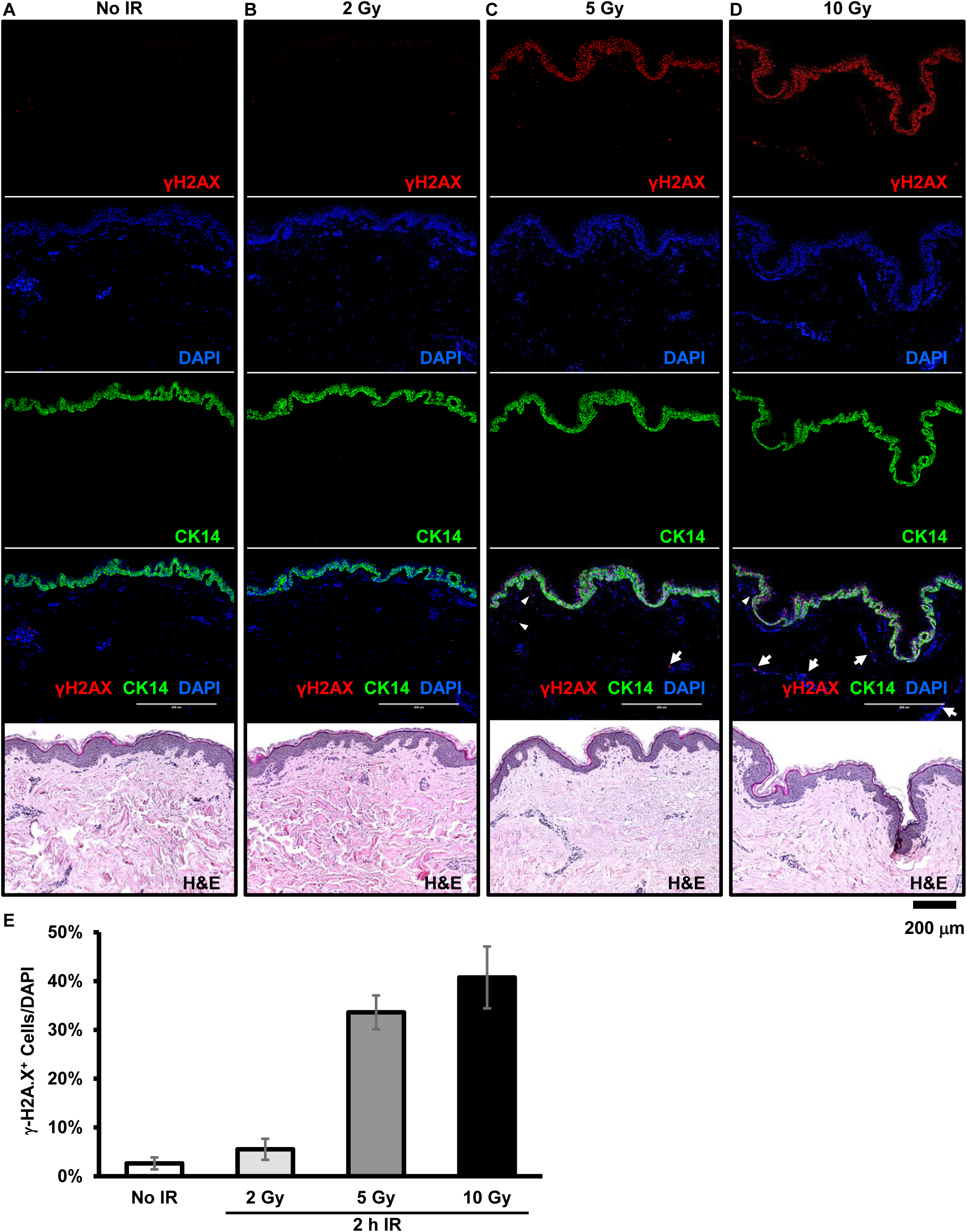
Dose-dependent DNA damage response in cutaneous tissues following *ex vivo* ionizing radiation (IR) exposure. (**A**) Representative H&E staining of control skin samples (no radiation). (**B**) Representative H&E staining of skin samples exposed to 2 Gy of ionizing radiation. (**C**) Representative H&E staining of skin samples exposed to 5 Gy of ionizing radiation. (**D**) Representative H&E staining of skin samples exposed to 10 Gy of ionizing radiation. Two hours post-IR, skin samples were collected and processed for immunohistochemical staining to visualize DNA damage (γH2AX, red), keratin 14 (CK14, green), and nuclei (DAPI, blue), alongside H&E staining (brightfield). (**E**) Quantification of DNA damage foci. Data are shown as mean ± SD (n = 3).

### X-ray irradiation induces genomic and mitochondrial DNA damage, leading to apoptosis in primary skin cells

DNA damage, in both the nuclear and mitochondrial genomes, affects numerous biological processes and disease pathways ^7^. To investigate the impact of ionizing radiation on DNA integrity, we examined nuclear DNA and mitochondrial DNA in primary skin cells, including keratinocytes, fibroblasts, and microvascular endothelial cells (MVECs), after exposure to 2, 5, 10, or 20 Gy of radiation. Long amplicon polymerase chain reaction (LA-PCR) revealed a dose-dependent loss of DNA integrity in both nuclear and mitochondrial genomes across all three cell types (Figure 2A, B).

**Figure 2.**
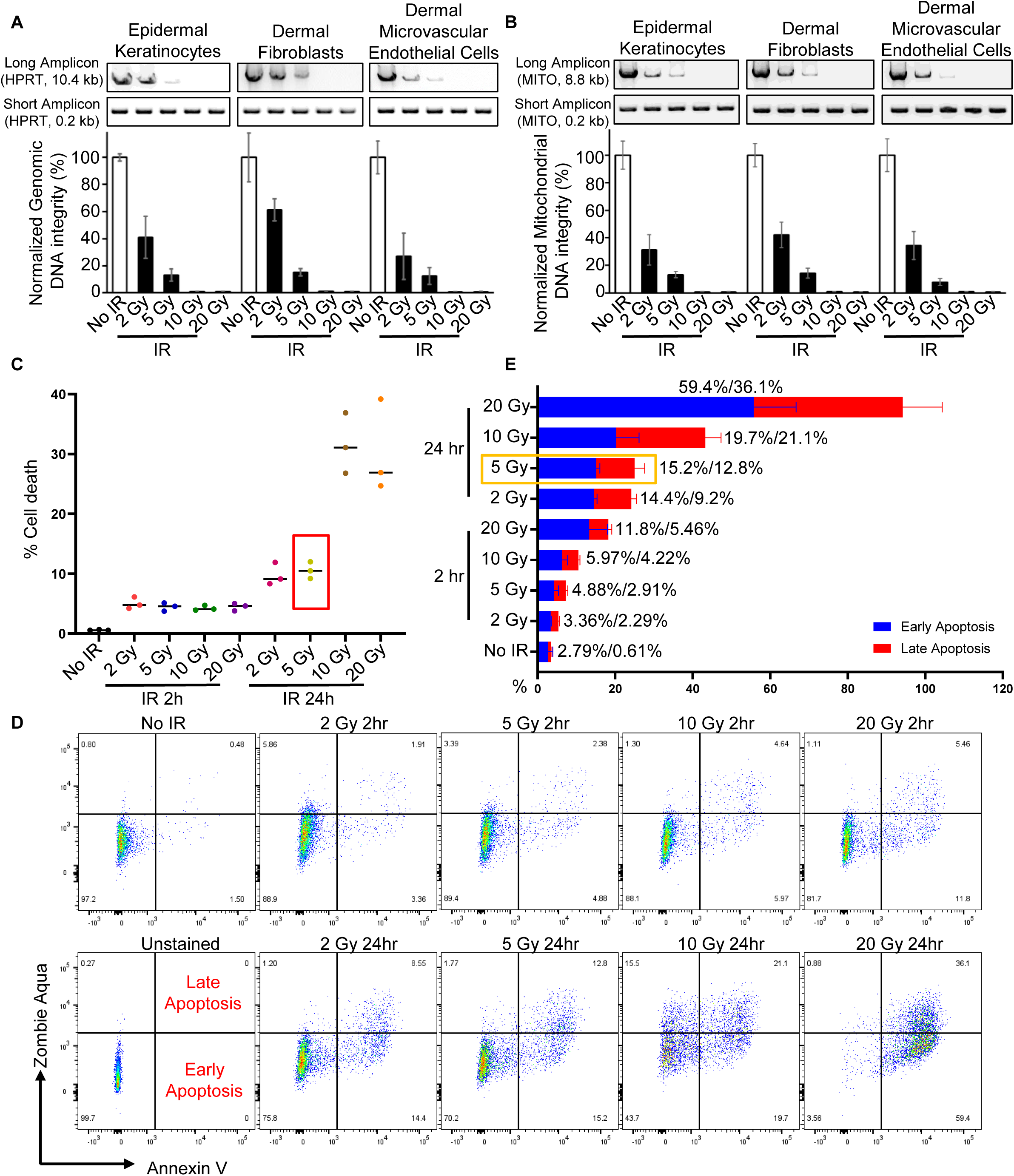
X-ray irradiation induces genomic and mitochondrial DNA damage, leading to apoptosis in primary skin cells. (**A-B**) Long amplicon PCR (LA-PCR) analysis of genomic and mitochondrial DNA integrity in human epidermal keratinocytes, dermal fibroblasts, and dermal microvascular endothelial cells immediately after exposure to different doses of X-ray irradiation (2 Gy, 5 Gy, 10 Gy, 20 Gy) or no irradiation (No IR, control). (**A**) Genomic DNA integrity was assessed using a 10.4 kb long amplicon from the HPRT gene and a 0.2 kb short amplicon from the same gene as a loading control. (**B**) Mitochondrial DNA integrity was assessed using an 8.8 kb long amplicon from the mitochondrial genome and a 0.2 kb short amplicon as a loading control. PCR products were resolved on agarose gels (0.8% for long amplicons; 2% for short amplicons). Representative gel images from three independent experiments are shown (top panels). Quantification of LA-PCR results normalized to the short amplicon controls is shown (bottom panels). (**C**) Quantification of cell death in human keratinocytes by flow cytometry at 2 hours and 24 hours post-exposure to the indicated doses of X-ray irradiation. (**D-E**) Representative flow cytometry plots and quantitative analysis of early and late apoptosis in human keratinocytes at 2 hours and 24 hours post-irradiation. Data are presented as mean ± SD (n = 3).

Since DNA damage can trigger apoptosis and cell death ^8–10^, we examined cell viability and apoptotic cell death in keratinocytes at 2-hour and 24-hour time points post-irradiation. Zombie Aqua and Annexin V were used to assess apoptosis: Zombie Aqua-positive cells indicate dead cells, Annexin V-positive cells indicate apoptotic cells, Zombie Aqua-negative/Annexin V-positive cells represent early apoptosis, and Zombie Aqua-positive/Annexin V-positive cells represent late apoptosis. Flow cytometry analysis indicated an increase in cell death proportional to both radiation dose and the time after exposure (Figure 2C). Further analysis showed that apoptosis accounted for 87% of the observed radiation-induced cell death (Supplementary Figure 1). Quantification of apoptotic cells at different radiation doses confirmed a dose-dependent increase in apoptosis (Figure 2D, E). Based on these findings, we selected 5 Gy irradiation as the optimal condition for subsequent experiments.

Collectively, our data demonstrate that ionizing radiation induces both nuclear and mitochondrial DNA damage, leading to apoptosis in a dose-dependent manner, with increasing manifestation of cellular damage over time.

### Telomerase enhances genomic DNA repair and promotes cell viability

Our previous study demonstrated that overexpression of telomerase reverse transcriptase (TERT) reduces DNA damage markers and extends lifespan in a murine model of premature aging ^5^. Therefore, we hypothesized that TERT expression might mitigate DNA damage and enhance cell viability following ionizing radiation. To test this hypothesis, we compared primary human aortic endothelial cells (HAECs) to HAECs with constitutive hTERT expression (Telo-HAECs) after exposure to 5 Gy of radiation. DNA damage signals, as assessed by immunofluorescence staining for γH2A.X, were reduced in Telo-HAECs compared to HAECs at 2- and 6-hours post-radiation (Figure 3A, B). Western blot analysis further confirmed reduced γH2A.X protein levels in Telo-HAECs compared to HAECs (Supplementary Figure 2A, B). Fewer apoptotic cells were observed in Telo-HAECs than in HAECs at various time points post-radiation (Supplementary Figure 2C). To confirm these findings, we induced DNA damage using bleomycin and observed a dose-dependent increase in DNA damage markers (γH2AX and 53BP1) and senescence-associated β-galactosidase activity in HAECs. In contrast, Telo-HAECs exhibited minimal induction of these markers (Supplementary Figure 2D–G), indicating that constitutive hTERT expression protects cells from both radiation- and bleomycin-induced DNA damage.

**Figure 3.**
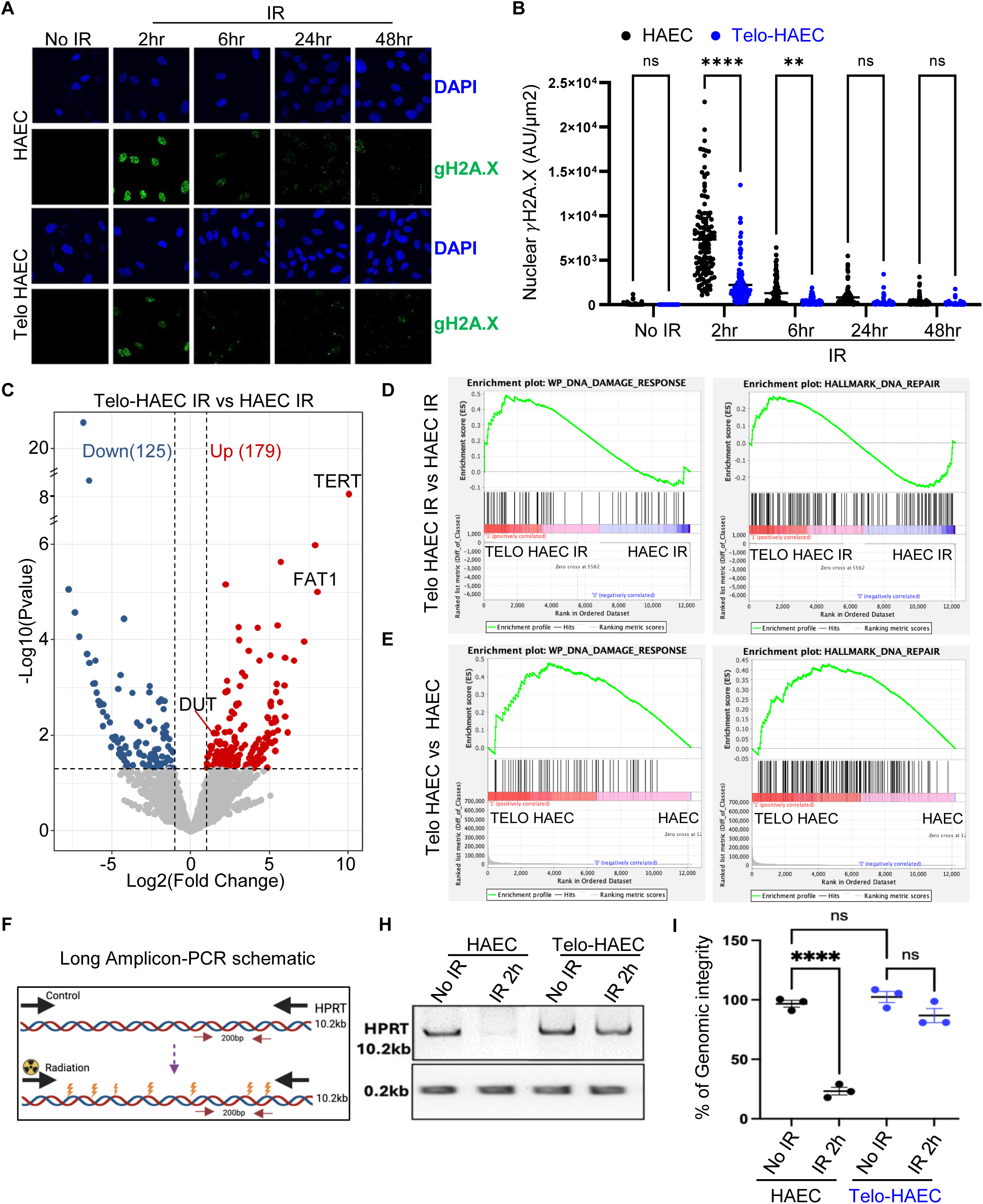
TERT expression in HAEC reduces DNA damage and increases cell survival. **(A, B**) Immunostaining for γH2AX and its quantification in HAEC and Telo-HAEC at different time points following 5 Gy irradiation, indicating DNA damage levels. (**C**) Volcano plots illustrating differentially expressed genes between Telo-HAEC and HAEC after irradiation exposure. A fold change of ≥ 2 and a p-value < 0.05 indicate significance. (**D-E**) Gene Set Enrichment Analysis (GSEA) demonstrates pathway enrichment to DNA damage response and DNA repair pathways in HAEC and Telo-HAEC, with or without irradiation. (**F**) Schematic representation of the Long Amplicon PCR (LA-PCR) assay used to assess DNA integrity. (**H, I**) Representative gel images and quantification of LA-PCR results from HAEC and Telo-HAEC under no irradiation (control) and 2 hours post-irradiation, highlighting differences in DNA damage and repair efficiency.

To explore the potential molecular mechanisms underlying TERT-mediated protection, we analyzed the transcriptional profile of HAECs and Telo-HAECs using bulk RNA sequencing, both with or without radiation. Upon radiation exposure, Telo-HAECs exhibited 179 upregulated and 125 downregulated genes compared to HAECs (Figure 3C, Supplementary Figure 2H–J and 3A). Gene Set Enrichment Analysis (GSEA) revealed enrichment of genes involved in DNA damage response and repair pathways (Figure 3D, Supplementary Figure 3B). Notably, even under basal conditions, Telo-HAECs exhibited differential gene expression profiles positively correlated with DNA repair pathways (Figure 3E), suggesting that TERT primes cells for DNA Damage response.

Given that TERT is classically associated with telomere maintenance, we assessed telomere length using two different methods Quantitative Fluorescent in situ (qFISH) and Terminal Restriction Fragment (TRF) analysis. Importantly, we found no significant differences in telomere length between irradiated and non-irradiated cells in both HAECs and Telo-HAECs (Supplementary Figure 3C, D). Nevertheless, the Telomeric Repeat Amplification Protocol (TRAP) assay confirmed the presence of telomerase activity in Telo-HAECs (but not HAECs) treated with bleomycin across various doses (Supplementary Figure 3E). Thus, the beneficial effect of telomerase expression is not due to telomere extension, but more likely related to a non-canonical effect of telomerase.

Additionally, in TERT expressing cells, transcriptomic analysis identified differential expression of genes involved in structural support, extracellular matrix organization, migration, cellular homeostasis, and stress responses, which are essential for cell survival and repair (Supplementary Figure 4D). Furthermore, the LA-PCR assays documented genomic DNA damage in all experimental groups; however, within 2 hours post-radiation, the genomic integrity of Telo-HAECs was restored, highlighting the accelerated DNA repair response of these cells (Figure 3F–I).

Together, these findings suggest that TERT enhances DNA repair and promotes cell viability independently of its canonical role in telomere elongation.

### TERT mRNA pretreatment enhances DNA repair to reduce radiation-induced apoptosis in primary skin cells

The recent success of mRNA-based therapies has highlighted the potential of lipid nanoparticles (LNPs) as a delivery system for mRNA. Our previous study suggested that TERT mRNA treatment reduces DNA damage in HGPS-derived iPSC fibroblasts and endothelial cells ^3–5^. Therefore, we hypothesized that cells could be protected from radiation-induced damage through pretreatment with TERT mRNA delivered via LNP. Since radiation-induced cell death is primarily mediated by apoptosis (Supplementary Figure 1B), we first examined whether TERT mRNA pretreatment could reduce radiation-induced apoptosis in keratinocytes, endothelial cells, and fibroblasts. Our previous work indicated that the DOTAP LNP formulation is efficient for *in vitro* transfection ^2^. Accordingly, we evaluated the efficiency of DOTAP LNP compared to commercially available RNA delivery reagents (Lipofectamine-Max and Jet-Messenger) in keratinocytes and fibroblasts using GFP mRNA. Confirming our prior work, DOTAP LNP exhibited the highest transfection efficiency (Figure 4A). Consequently, we used DOTAP to deliver TERT mRNA for subsequent experiments. Cells were pretreated with vehicle, GFP mRNA, or TERT mRNA 24 hours prior to 5 Gy radiation exposure, and apoptotic cells were quantified 24 hours post-radiation (Figure 4B). Pretreatment with TERT mRNA reduced the apoptotic cell population, including both early and late apoptosis, in keratinocytes and MVECs compared to vehicle or GFP mRNA (Figure 4C–I). A similar trend, though not statistically significant, was observed in fibroblasts (Supplementary Figure 5), suggesting cell type-specific variability in response to TERT mRNA pretreatment.

**Figure 4.**
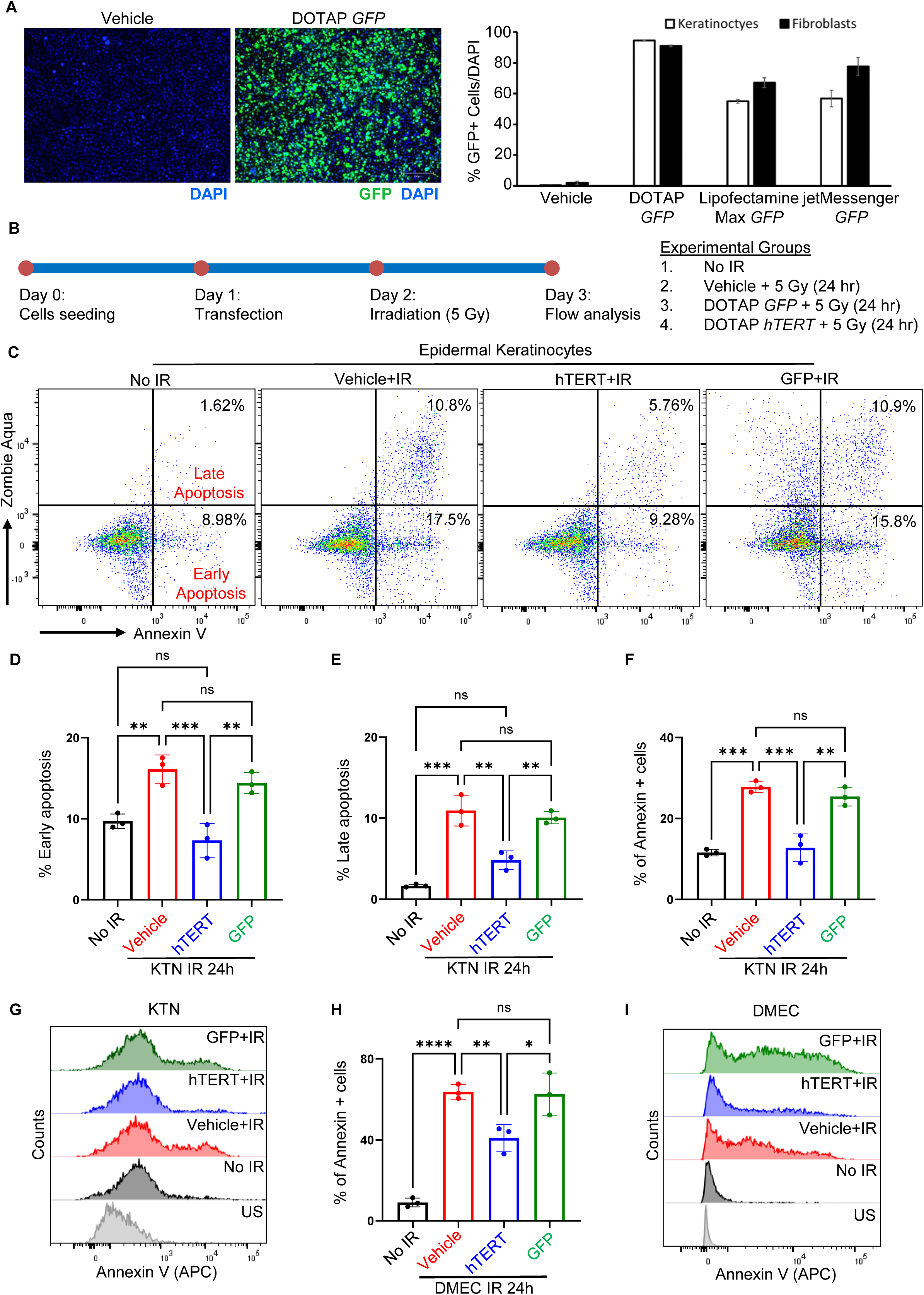
TERT mRNA treatment reduces radiation-induced apoptosis in primary skin cells. (**A**) Human epidermal fibroblast and keratinocytes were treated with vehicle control or GFP mRNA encapsulated by cationic lipid nanoparticle DOTAP or mixed with commercial transfection reagents (Lipofectamine Max or jetMessenger) to assess transfection efficiency. Representative images of green fluorescence from live cells (left panel) and DAPI-stained nuclei are shown (n = 3 independent experiments). Quantification GFP+ cells/DAPI are presented as mean percentage ± SD (n = 3). (**B**) Schematic outline of the experimental setup. (**C-E**) Representative flow cytometry plots and quantitative analysis of early and late apoptosis in human epidermal keratinocytes, assessed 24 hours after ionizing radiation (5 Gy) or no irradiation (No IR control). (**F, G**) Representative flow cytometry plots and quantitative analysis of Annexin V-positive cells in indicated groups of human epidermal keratinocytes. (**H, I**) Representative flow cytometry plots and quantitative analysis of Annexin V-positive cells in human dermal microvascular endothelial cells treated with indicated groups. Primary cells were transfected with DOTAP-encapsulated GFP or hTERT mRNA or treated with vehicle control. Flow cytometry analysis was performed 24 hours following 5 Gy irradiation or No IR control. Early apoptosis was identified as Zombie Aqua-negative and Annexin V-positive cells, while late apoptosis was identified as Zombie Aqua-positive and Annexin V-positive cells. Data are shown as mean ± SD (n = 3). ns, *P* > 0.05; *, *P* < 0.05; **, *P* < 0.01; ***, *P* < 0.001, ****, *P* < 0.0001. *P* values were calculated using two-way ANOVA. KTN: Epidermal Keratinocytes; DMEC: Dermal Microvascular Endothelial Cells.

To investigate the potential mechanism of TERT mRNA treatment in reducing apoptosis, we examined genomic DNA integrity in keratinocytes, fibroblasts, and MVECs following 5 Gy radiation at various time points (0, 0.5, 1, 2, 6, and 24 hours). LA-PCR analysis revealed progressive loss of DNA integrity over time, which was mitigated by TERT mRNA pretreatment in all cell types (Figure 5A–C). These findings are consistent with the protective effects observed in Telo-HAECs (Figure 3), suggesting that TERT mRNA pretreatment enhances DNA repair.

**Figure 5.**
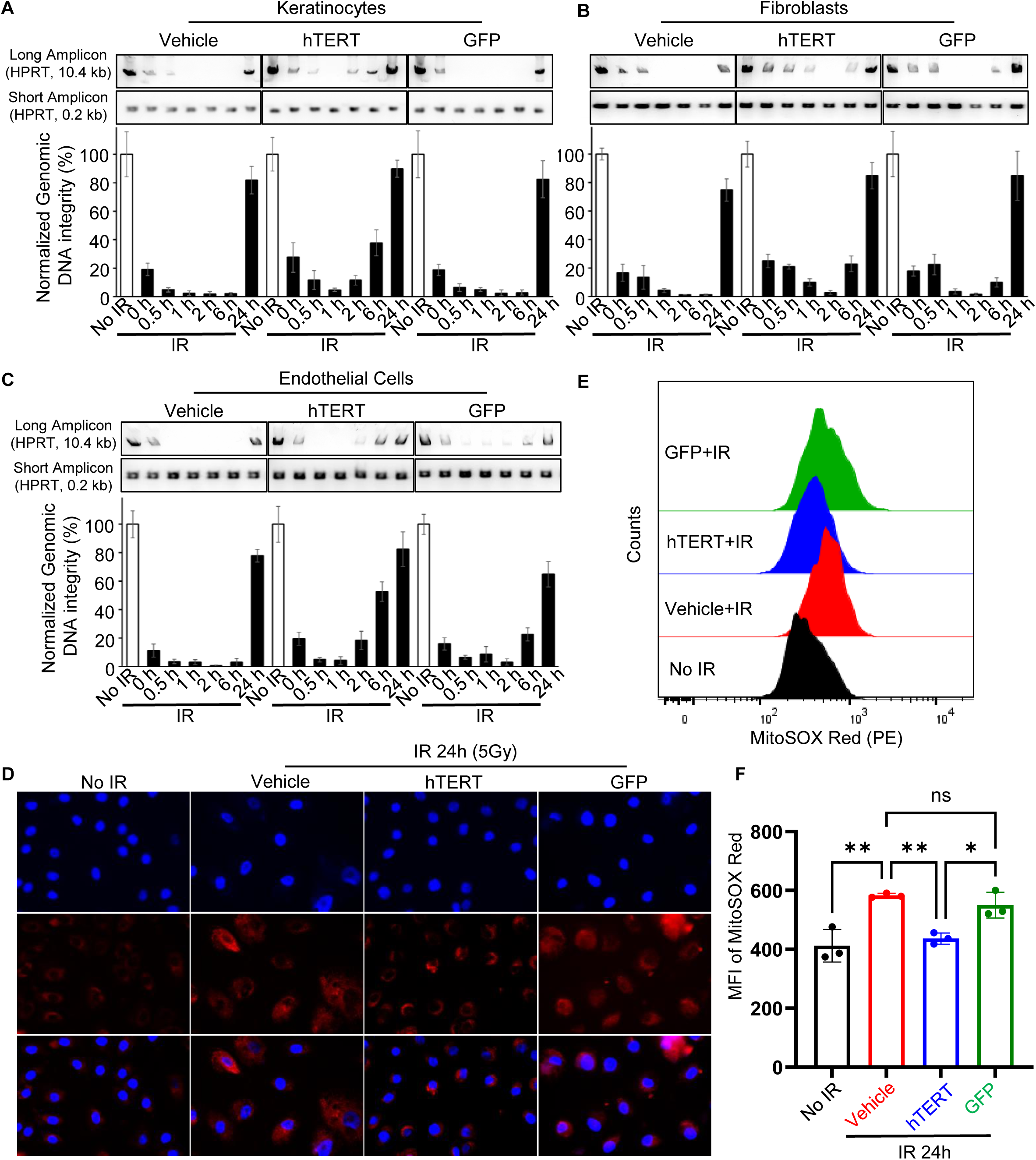
Telomerase mRNA treatment accelerates repair in radiation-induced DNA damage. **(A-C**) Quantification of DNA damage by long amplicon PCR (LA-PCR) analysis of genomic DNA isolated from human epidermal keratinocytes, dermal fibroblasts, and dermal microvascular endothelial cells exposed to 5 Gy irradiation or no irradiation. At 0 h, 0.5 h, 1 h, 2 h, 6 h, and 24 h post-irradiation, DNA was analyzed using primers for a long amplicon (10.4 kb region in the HPRT gene) and a short amplicon (0.2 kb region in the HPRT gene) as a loading control. PCR products were resolved on agarose gels (0.8% for long amplicons, 2% for short amplicons). Representative gel images from three independent experiments are shown (top panels), with quantification of genomic DNA integrity normalized to the short amplicon control (bottom panels). (**D**) Representative images of MitoSOX analysis at 24 hours post-irradiation in keratinocytes treated with vehicle control, GFP mRNA, or TERT mRNA. (**E-F**) Representative flow cytometry plots and quantitative analysis of MitoSOX Red fluorescence in keratinocytes treated with vehicle, GFP mRNA, or TERT mRNA 24 hours after irradiation, indicating differences in mitochondrial superoxide production. MFI: Mean Fluorescence Intensity.

Telomerase protects against oxidative stress by reducing mitochondrial reactive oxygen species (ROS) production and preventing cellular apoptosis ^11,12^. Elevated ROS levels are a hallmark of mitochondrial damage and dysfunction ^13,14^. We next investigated whether TERT mRNA pretreatment reduces mitochondrial ROS levels. Using the MitoSOX assay, we observed lower mitochondrial ROS levels in TERT mRNA-pretreated keratinocytes when compared to vehicle or GFP-treated cells at 24 hours post-radiation (Figure 5D–F). These data suggest that TERT mRNA pretreatment enhances the repair of both genomic and mitochondrial DNA damage to reduce radiation-induced apoptosis.

### Optimal delivery system of TERT mRNA pretreatment for the human skin

To investigate the clinical applicability of TERT mRNA pretreatment, we optimized its delivery using *ex-vivo* human skin tissue. We tested various LNP formulations (MC3, DOTAP, and JetMessenger) in combination with microneedling. Among the tested formulations, MC3 LNP exhibited the highest transfection efficiency (Figure 6A–H). In addition, the TRAP assay confirmed that the delivered TERT mRNA generated active TERT protein in MC3-treated skin 24 hours post-delivery, compared to GFP-treated skin. (Figure 6I). These results indicate that MC3 LNP-mediated delivery of TERT mRNA using microneedling is an effective approach for human skin.

**Figure 6.**
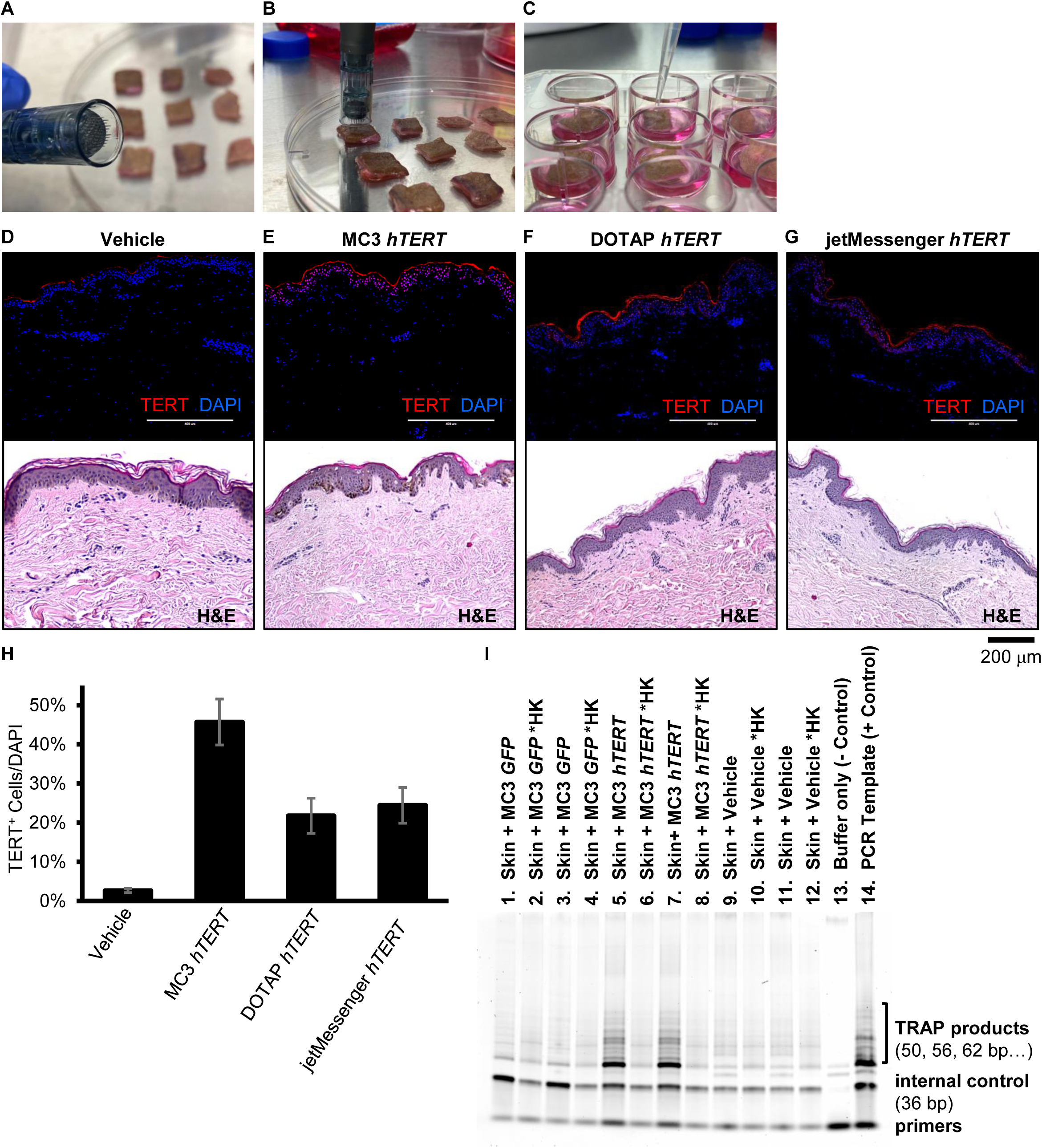
Optimization of mRNA delivery to human skin explants. (**A, B**) 1-cm2 pieces of human skin underwent microneedling using a 36-pin microneedling pen. (**C**) Following the procedure (6 repetitions per second for 20 s, 2.0 mm depth), skin explants were treated with hTERT mRNA and cultured at air-liquid interface. (**D-G**) Immunofluorescence and H&E staining of ex-vivo skin samples subjected to microneedle delivery of hTERT mRNA carried by ionizable lipid nanoparticle MC3, cationic lipid nanoparticle DOTAP, or commercial transfection reagent jetMessenger. Vehicle solution served as the control for transfection. (**H**) Quantification of TERT+ cells (red) over nuclei (DAPI, blue) are presented in graph. Data are shown as mean ± SD (n = 3). * p < 0.05 which were analyzed using two-way ANOVA. (**I**) TRAP assay for telomerase activity in the ex vivo cutaneous tissues following microneedle delivery of MC3 encapsulated mRNA or vehicle control. Lysates from skin samples treated with MC3 lipid nanoparticle encapsulated GFP (lanes 01-04), hTERT (lanes 05-08), or vehicle control (lanes 09-12). Lysis buffer served as negative control (lane 13) and PCR amplified template was used as a positive control (lane 14). Even number lanes were loaded with heat-killed (HK; 85 °C for 10 min to inactivate telomerase) lysates set up in parallel for each sample extract.

### Pretreatment with telomerase mRNA reduces radiation-induced DNA damage and apoptosis in human skin

Next, we investigated whether TERT mRNA pretreatment reduces radiation-induced DNA damage and enhances cellular survival in *ex vivo* human skin. Human skin samples were pretreated with vehicle, GFP mRNA, or TERT mRNA via microneedling, incubated for 24 hours, and subsequently exposed to 5 Gy radiation. To assess DNA damage, we quantified γH2AX immunofluorescent staining. Sequential H&E staining was performed in parallel to indicate the analyzed area and signal localization within the epidermis. Radiation induced an substantial increase in γH2AX in skin samples treated with vehicle or GFP, but not in TERT-treated skin (Figure 7A, B). These findings were consistent with the results obtained from cultured HAECs (Figure 3A, B). Radiation induced an increase in p16 and p21, which was attenuated in TERT-pretreated skin (Figure 7C).

**Figure 7.**
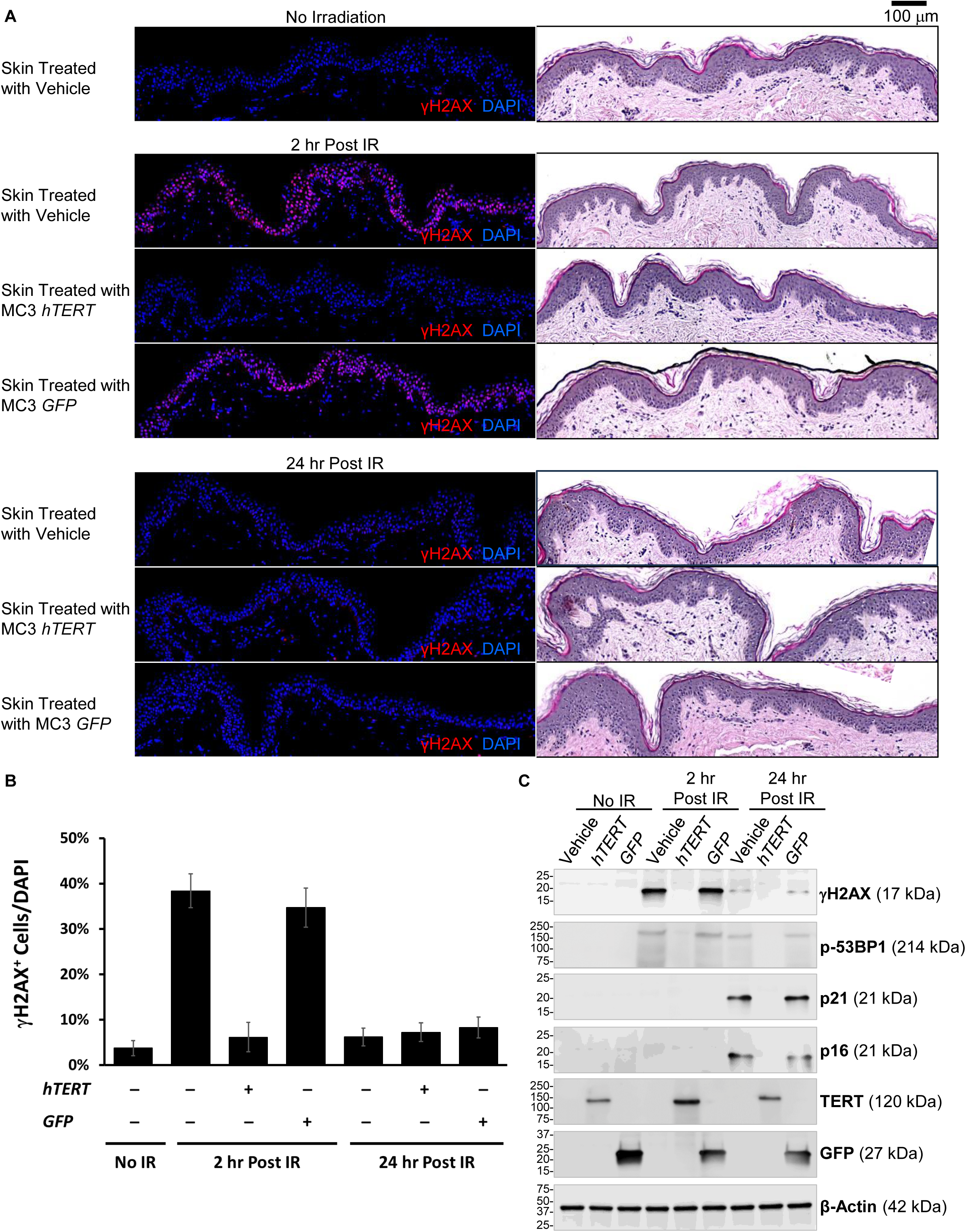
Treatment of ex-vivo skin with hTERT mRNA via microneedling reduced radiation-induced γH2AX and senescence markers. (**A**) Human skin (1-cm2 pieces) undergone microneedling procedure (6 repetition per second for 20 s, 2.0 mm depth) were immediately applied with either vehicle solution or encapsulated mRNA (hTERT or GFP) within ionizable lipid nanoparticle MC3. Following 24 hr of ex-vivo culture, skin explants were exposed to X-ray irradiation (5 Gy) and harvested 2 hr or 24 hr post irradiation for immunohistochemical staining (γH2AX, red; nuclear DAPI, blue; H&E, brightfield). A group of skin samples treated with vehicle and unexposed to radiation (no irradiation) served as control. Representative images shown, from 3 independent experiments. (**B**) Summary data showing the percentage of cells stained positive for DNA damage marker γH2AX. Data are shown as mean ± SD (n = 3). * p < 0.05 which were analyzed using two-way ANOVA. (**C**) Immunoblot analysis of whole cell lysates collected from non-irradiated skin samples treated with vehicle (lane 1), hTERT (lane 2), GFP (lane 3), or from skin samples harvested 2 hr (lane 4-6) or 24 hr (lane 7-9) post irradiation (5 Gy). β-actin antibody was used as a loading control.

To evaluate apoptosis, we performed TUNEL (terminal deoxynucleotidyl transferase-mediated dUTP nick end labeling) assay on irradiated human skin samples. TERT-pretreated skin exhibited significantly fewer apoptotic cells compared to vehicle- or GFP-pretreated skin at 24 hours post-radiation (Figure 8A, B). To confirm that the protective effect of pretreated TERT mRNA was independent of telomere function, we measured telomere length before and after radiation in all samples. No changes in telomere length were detected with qFISH (Figure 8C). Apurinic/apyrimidinic (AP) sites occur when DNA bases are lost due to damage from reactive oxygen species (ROS), radiation, alkylating agents, or spontaneous hydrolysis, and represent one of the most common types of DNA damage. We quantified AP sites in DNA isolated from skin samples pretreated with vehicle, GFP, and TERT mRNA at 24 hours post-radiation. A significant number of AP sites were present in DNA extracted from vehicle- or GFP-pretreated skin, while TERT-pretreated skin exhibited a marked reduction in AP sites at 24 hours post-radiation, confirming enhanced DNA repair (Figure 8D, E). Collectively, these findings demonstrate that TERT mRNA pretreatment enhances DNA repair and reduces apoptosis in human skin following radiation exposure.

**Figure 8.**
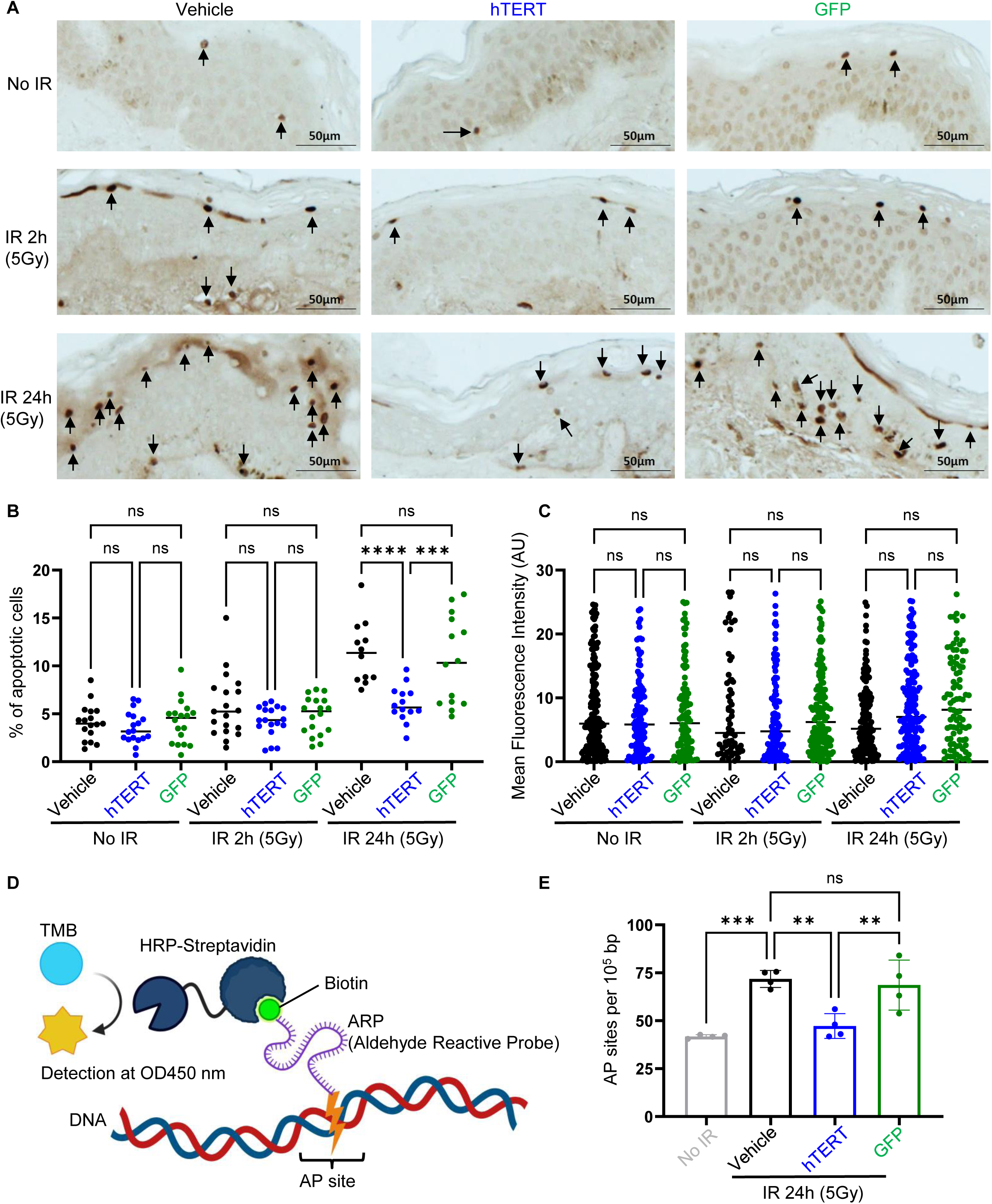
TERT mRNA treated skin reduces apoptotic cells due to enhanced DNA repair. (**A**) Representative figures of skin samples with TUNEL staining. (**B**) The quantification of the TUNEL assay indicates the percentage of apoptotic cells in skin samples treated with TERT or GFP mRNA or vehicle. (**C**) qFISH analysis of skin samples treated with TERT or GFP mRNA or vehicle. (**D**) The schematic graph illustrates the colorimetric assay for detecting DNA damage (AP sites). (**E**) Colorimetric analysis of AP sites from DNA extracted from skin samples treated with TERT or GFP mRNA or vehicle. Data are shown as mean ± SD (n = 3-5). ** *P* < 0.01, *** *P* < 0.001, **** *P* < 0.0001, which were analyzed using one-way ANOVA. TMB: 3,3’,5,5’-Tetramethylbenzidine.

## Discussion

In the current study we observed that prior treatment with mRNA hTERT substantially reduced radiation induced genomic and mitochondrial DNA damage and apoptosis in human skin cells. The effect of TERT mRNA pretreatment enhances DNA repair, reduces mitochondrial ROS, and decreases apoptosis without extending telomere length during the experimental period suggesting a non-canonical function of TERT to accelerate the cellular recovery from radiation. These findings highlight a potential therapeutic approach for preventing radiation-induced skin injury.

Analysis of the transcriptome profile of constitutive TERT expression in HAECs (Telo-HAEC) has revealed higher basal levels of genes that could equip the cells to handle cellular stress, including the response to radiation-induced damage ^14^. After radiation, transcriptional profiling revealed that in Telo-HAEC, there was differential upregulation of pathways associated with cell migration, angiogenesis, wound healing, and tissue generation, such as NABA ECM glycoproteins pathway (Networks of ECM (Extracellular Matrix)-Associated proteins), or Hallmark of Epithelial-Mesenchymal Transition. Furthermore, the downregulation in Telo-HAEC of pathways associated withcell junction organization, would also favor increased cell migration and wound healing processes. While the expression of TERT has been primarily studied for its role in the context of cancer, our data suggest that it may also prime cells against specific stress responses, particularly when analyzing pathway enrichment for both DNA damage response and repair. Genes in these two pathways are upregulated in Telo-HAEC, which could account for the strong recovery in DNA repair observed shortly after radiation exposure. Others have suggested the involvement of TERT in DNA repair ^15–17^, although its direct role is unclear; however, our data suggest it operates through a non-telomeric mechanism possibly mediated by gene regulation. Importantly, we observed that enhanced cellular viability and DNA repair do not require the constitutive expression of TERT, since transient expression of TERT using mRNA, in both *ex-vivo* skin and keratinocytes, reduces γH2A.X, AP sites, and apoptosis. Interestingly, TERT expression in keratinocytes reduced mitochondrial ROS, which was significantly high after radiation exposure in untreated cells, suggesting a key process in the recovery of cellular function by TERT ^18^. Importantly, when TERT mRNA was delivered using lipid nanoparticles to human *ex-vivo* skin, the beneficial effects observed *in vitro* were recapitulated in a model closer to clinical application. Importantly, two hours post-radiation, a significant reduction of γH2A.X foci was observed using immunostaining. Indeed, the long amplicon PCR (LA-PCR) and AP (apurinic/apyrimidinic) site analysis indicate enhanced DNA repair in both *in vivo* and *in vitro*. The transcriptional response to radiation was different in Telo-HAEC versus HAEC.

Accumulation of DNA damage is a key factor in cellular aging. Damage to genomic as well as telomeric DNA can induce gene dysregulation, and activation of DNA repair pathways. DNA damage generates ROS and drives chronic inflammation and cellular senescence. Cells have evolved sophisticated mechanisms to manage various types of DNA damage. When exposed to harmful agents like pollution or radiation, cells halt division by upregulating checkpoint proteins, allowing them to repair DNA and maintain genomic integrity. While some cells successfully repair damage and continue functioning, others face an overwhelming challenge, especially from double-strand breaks, which can lead to apoptosis. Cells that partially resolve damage may enter senescence. Senescent cells actively release inflammatory factors, a state known as the senescence associate secretory phenotype (SASP). Sustained non-lethal damage, such as low-grade radiation or chemotherapy, can prematurely induce senescence, leading to chronic inflammation and oxidative stress.

Recent molecular and cellular studies indicate that the damage to normal tissues following radiation may be due to ROS and reactive nitrogen species (RNS), as well as pro-inflammatory cytokines and chemokines, which alter or degrade tissue and organ function ^19–21^. A major source of chronic ROS and inflammation is radiation-induced senescent cells, as evidenced by biomarkers such as DNA damage response pathways, mitochondrial dysfunction, and oncogene activation ^22–25^. Radiation-induced senescence in mouse skin ^26^ and various types of cultured cells have been demonstrated ^27,28^. Importantly, the elimination of senescent cells by senolytic drugs has been shown to improve the recovery from muscle weakness and pulmonary fibrosis induced by radiation ^29,30^. Several strategies to block proinflammatory molecules or reduce oxidative stress have been shown to prevent or alleviate radiation toxicity. For instance, pan-suppression of macrophage infiltration and cytokines/chemokines has been observed to mitigate effects in normal tissues, including the skin and brain ^31–33^.

Our data indicates that enhancing the repair of genomic DNA damage with TERT mRNA therapy increases cellular viability and reduces markers of senescence in ex-vivo skin and various cell types exposed to radiation. While TERT is commonly associated with telomere extension, our data show that its role in DNA repair extends beyond telomeres. Notably, the reduction in DNA damage marker γH2A.X occurs within 2 hours, suggesting no telomere extension is involved, as telomere elongation is replication dependent. Consistently, qFISH analysis revealed no significant changes in telomere length at 2, 24, or even 72 hours post-radiation or post-bleomycin treatment. Telomere shortening after radiation depends on dose, frequency, and cell type, and 72 hours may not be sufficient for significant changes in telomeres in endothelial cells, given their limited divisions during this time. Preclinical models and patients with genetic causes of defective DNA repair are more radiosensitive, as in Fanconi Anemia (FA), Nijmegan Break Syndrome, Bloom’s Syndrome, and Ataxia Telangiectasia ^34^. These studies suggest that enhancing DNA damage repair could preserve the homeostasis of the cells/tissue that are not the target of the therapy.

DNA damage could induce genomic instability and enhance the risk of mutations in tumor suppressor genes, leading to cancer. While we and others have shown that TERT expression in normal cells does not initiate cancer, telomerase becomes activated in 80% of cancer cells. This activation not only extends their telomeres, providing greater proliferation capabilities, but also reduces DNA damage, and increases resistance to radiation therapy. Our findings offer valuable insight into the pathological effects of TERT expression in cancer cells, beyond its role in telomere maintenance.

Previously, we have demonstrated that treatment with hTERT-mRNA restores telomere length, reduces senescence-associated secretory phenotype (SASP), and DNA damage, and rescues normal cellular functions in endothelial and vascular smooth muscle cells derived from patients with Hutchison Gilford Progeria syndrome (HGPS) ^5^. Furthermore TERT overexpression in a HGPS mouse model reduced DNA damage markers and inflammatory cytokines in the systemic vasculature and extended the lifespan of the animals. These observations may not be entirely explained by the canonical function of telomerase.

The current findings highlight that pretreatment with TERT enhances cellular repair following radiation exposure, suggesting its potential as a therapeutic strategy to improve the safety and effectiveness of treatments for radiation-damaged normal skin.

## Material and methods

### Messenger RNA Synthesis

Human *TERT* (NM_198253.2) and *GFP* mRNA were synthesized by Houston Methodist Research Institute (HMRI) RNA Core by in vitro transcription (IVT) with pseudouridine added to the nucleotide mix to reduce inflammatory activation and to improve translation, as previously described (Ramunas et al., 2015, FASEB J). mRNA was purified by filtration and assessed for integrity using a TapeStation (Agilent).

### Lipid Nanoparticle (mRNA LNP) Formulation

Due to the chemical and enzymatic instability of naked mRNA in physiological fluids, *in vivo* mRNA administration requires a delivery system that protects mRNA from degradation and effectively transfects the target cells. Lipid nanoparticles (LNP) are often used for mRNA delivery, as in the SARS CoV 2 mRNA vaccines. In previous studies, we employed two types of lipid nanoparticles (LNPs), MC3-LNP and DOTAP, to optimize TERT mRNA delivery in ex vivo skin models and primary cells.

In the current study, mRNA LNP was formulated using a molar ratio of DSPC, DMG-PEG2000, cholesterol, and Dlin-MC3-DMA at 8:1.5:38.5:52 respectively. DOTAP mRNA LNP was prepared with a molar ratio of DOPE, DOTAP, and DMG-PEG2000 at 49:49:2 respectively. The lipid components were dissolved in ethanol, while mRNA was dissolved in citric buffer (100 mM, pH 5.0). Mixing occurred at a flow rate of 3:1 aqueous to ethanol, with a total flow rate of 10 mL/min, followed by dialysis in PBS at 4°C for a minimum of 8 hours. The mRNA LNP were formulated using the NanoAssemblr Benchtop (Precision Nanosystems).

### Cell Culture

Human adult epidermal keratinocytes (Lonza #00192627) were grown in KGM-Gold medium (Lonza #00192060). Human adult human dermal fibroblasts (Lonza #CC-2511) were cultured in FGM medium (Lonza #CC-3132). Human dermal microvascular endothelial cells (Lonza #CC-2516) were grown in EGM2 medium (Lonza #CC-3162). Human Primary Aortic Endothelial Cells (HAEC, ATCC # PCS-100-011) and TeloHAEC (ATCC # CRL-4052) were grown in EGM2 medium. Cells were maintained at 37□C and 5% CO2.

### *EX-Vivo* culture of human skin

De-identified adult human skin samples were collected under the Houston Methodist Research Institute Institutional Review Board approved protocol with patient/family consent (IRB #PR000027413). Hypodermal fat layer of the skin was removed, then cut into 1-cm^2^ pieces using disposable scalpel (Integra Miltex #4-410). Skin explants were cultured in keratinocyte growth medium (ATCC #PCS-200-040) at 37°C, 5% CO2, maintaining an air-liquid interphase as described (Corzo-León et al., 2019, Methods).

### Transfection with mRNA Lipid Nanoparticles

Primary human skin cells were seeded 1 x 10^5^ cells/well in 12-well culture plate or 2.5 x 10^5^ cells/well in 6-well culture plate. DOTAP mRNA LNP (*hTERT* or *GFP*) was added 1 μg/mL of culture media. Human skin explants were subjected to microneedling by Dr. Pen Ultima M8 microneedling device, 36 pin, 6 repetition per second for 20 s, 2.0 mm depth.

### Histological Analysis

Skin explant was fixed in 10% formalin, paraffin embedded, and sectioned (5 μ by the HMRI Pathology Core Laboratory. Histological section was stained with hematoxylin and eosin (H&E) for tissue morphology or immunostained using anti-γH2AX antibody (Abcam #ab81299), anti-CK14 antibody (Abcam #ab7800) or anti-TERT antibody (Novus #NBP3-14565). Slides were preserved with VECTASHIELD HardSet Antifade Mounting Medium with DAPI (Vector Labs #H-1500-10) and imaged by EVOS-FL-Auto-Imaging System (Life Technologies).

### Western Blot Analysis

Total cell lysates were collected from skin explants (n = 3) using handheld rotor-stator homogenizer (Qiagen TissueRuptor II #9002755) and T-Per protein extraction reagent (ThermoFisher #78510) according to manufacturer’s protocol. Whole cell lysates (25 ng/lane) were resolved by SDS/PAGE (Bio-Rad Any kD Mini Gel #4569036) and transferred to PVDF membrane (Bio-Rad #1704156), followed by immunoblotting using SuperSignal West Pico PLUS chemiluminescent reagent (ThermoFisher #34580) and imaged by ChemiDoc MP Imaging System (Bio-Rad #12003154).

### Telomerase Activity Detection

For Telomeric Repeat Amplification Protocol (TRAP) assay (MilliporeSigma TRAPeze Telomerase Detection Kit #S7700) total cell lysates were collected from skin explants (n = 3) using handheld rotor-stator homogenizer (Qiagen TissueRuptor II #9002755) and CHAS lysis buffer according to manufacturer’s protocol. Samples were diluted to equal protein concentration (0.03 µg/µL). For heat-kill (HK) control, each sample was subjected to 85°C for 10 minutes to inactivate telomerase. Following 32 cycles of PCR amplification, TRAP products were resolved by gel electrophoresis. Quantification of telomerase activity in each sample was normalized to HK control.

### MitoSOX detection assay

The MitoSOX (Cat# M36008) reagent was used to detect mitochondrial ROS according to the manufacturer’s protocol. In brief, the 5 mM MitoSOX reagent stock solution was diluted in HBSS buffer (provided by the kit) to make a 5 µM solution. Cells prepared in a 24-well plate were incubated with the 5 µM MitoSOX reagent working solution for 10 minutes at 37°C, protected from light. Following incubation, the cells were washed with PBS and either imaged using a fluorescence microscope or collected for ROS quantification via flow cytometry.

### Apoptosis analysis

For cell apoptosis analysis, cells were collected and stained using the Zombie Aqua Fixable Viability Kit (BioLegend) according to the manufacturer’s instructions. After staining, the cells were washed with PBS, resuspended in Annexin V binding buffer (catalog number 422201; BioLegend), and incubated with APC Annexin V (catalog number 640920; BioLegend) for 15 minutes at room temperature in the dark. The samples were then analyzed using an LSR II or Fortessa flow cytometer. Data analysis was performed using FlowJo software, version 10 (Tree Star, Inc., Ashland, OR, USA).

### TUNEL assay

Human skin tissue from four donors was used for the TUNEL assay (Abcam ab206386), conducted according to the manufacturer’s instructions. Paraffin-embedded tissue samples were cut into sections and mounted on glass slides, followed by rehydration through sequential immersion in Xylene Substitute (Safeclear II, Fisher HealthCare) and graded ethanol solutions. The slides were washed with 1x TBS, permeabilized with proteinase K solution, and treated with Hydrogen Peroxide to quench endogenous peroxidase activity. The tissues were labeled with Terminal deoxynucleotidyl transferase) and developed using DAB (3,3′-Diaminobenzidine) solution. Total cells were stained with Methyl Green, and specimens were mounted with mounting media. Images were captured using an Olympus BX61 microscope at 40x magnification. Apoptotic cells in each specimen were counted in at least five fields of view, ensuring a minimum of 150 cells per field, and data were analyzed using GraphPad Prism 10.

### qFISH analysis

Cells or cut sections from tissues were fixed in 3.7% formaldehyde, followed by dehydration with an ethanol series: 5 minutes each in 70%, 85%, and 100%. The telomere probe (PNA Bio F1013 TelC-Alexa 647) was added to the slide and incubated overnight at 4°C. After washing twice with 70% formamide and twice with 1% BAS respectively, the slides were dehydrated with ethanol. DAPI was added, and telomere length was reported as the intensity of probe normalized to the area of nuclei.

### Beta-gal analysis

Beta-galactosidase staining was performed on cultured cells following the manufacturer’s protocol (Senescence β Galactosidase Staining Kit, Cell Signaling #9860). In brief, cells were fixed in the plate and washed with PBS, followed by incubation with a staining solution containing X-gal (5-bromo-4-chloro-indoxyl) in citrate-phosphate buffer (pH 6.0). Blue-stained cells were visualized under a microscope to identify senescent cells. The positive cells were quantified using ImageJ software.

### Long-Amplicon Polymerase Chain Reaction (LA-PCR)

DNA was isolated using a DNeasy Blood and Tissue Kit (Qiagen #69504) according to the manufacturer’s instructions. LA-PCR was performed from a modified protocol previously described (Dharmalingam et al., 2020, ACS Nano). Briefly, 10 ng DNA from each sample was amplified with LongAmp Taq DNA Polymerase (NEB #M0323) for genomic and mitochondrial DNA amplicons, using the primer sequences and PCR conditions given in Supplemental Table 1. PCR products were separated by gel electrophoresis (0.8% agarose gel for long amplicon products and 2% agarose gels for short amplicon products), stained by GelRed nucleic acid stain (MilliporeSigma #SCT123), and imaged by ChemiDoc MP Imaging System (Bio-Rad #12003154). Quantification of PCR products was based on band intensity using ImageJ software.

### RNA sequencing data analysis

Low-quality reads were removed, and the remaining high-quality sequences were aligned to the human genome reference (GRCh38) using the STAR aligner. Differential gene expression analysis was performed using the DESeq2 package. Genes with an absolute fold change 2 ≥ and an adjusted p-value (p-adj) < 0.05 were considered significantly differentially expressed genes (DEGs). Pathway analysis of DEGs was conducted using Gene Ontology (GO) term analysis. Additionally, Gene Set Enrichment Analysis (GSEA) of DNA damage and DNA repair pathways was performed using the GSEA R package.

### Data analysis

Data visualization and statistical analyses were performed using R and GraphPad Prism. Measurement data are presented as Mean ± SD. Comparisons between two groups were conducted using Student’s t-test, while comparisons among multiple groups were carried out using one-way or two-way ANOVA.

**Table 1.**
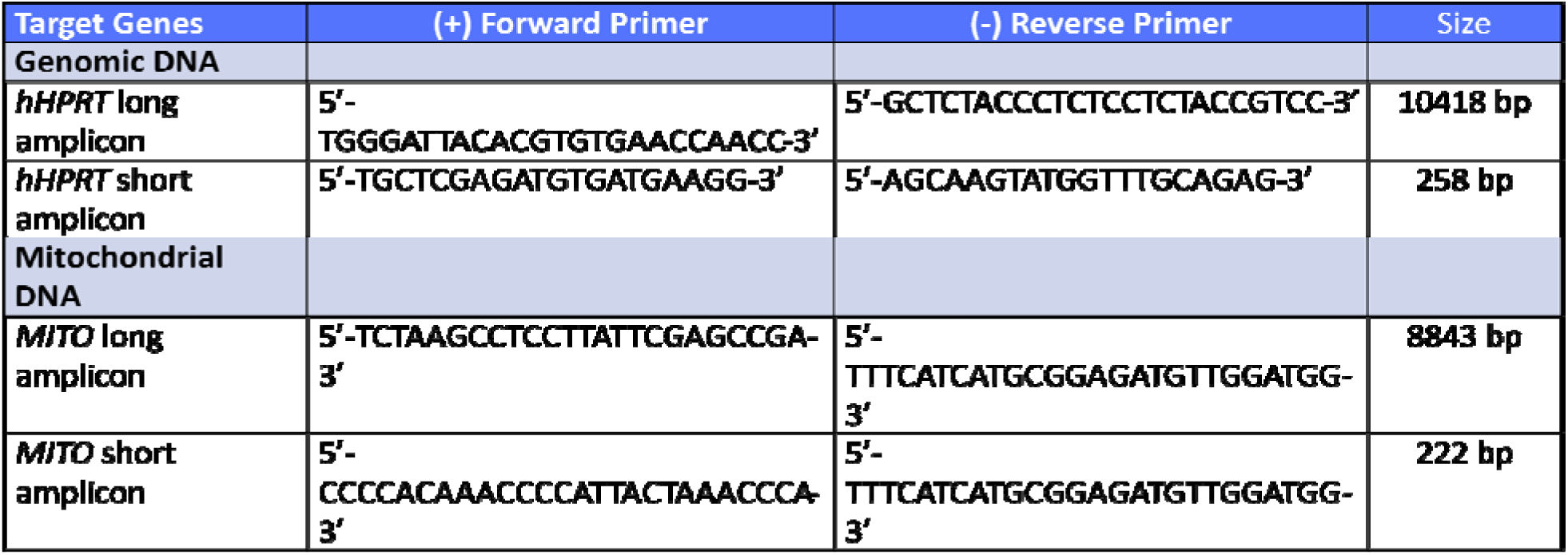
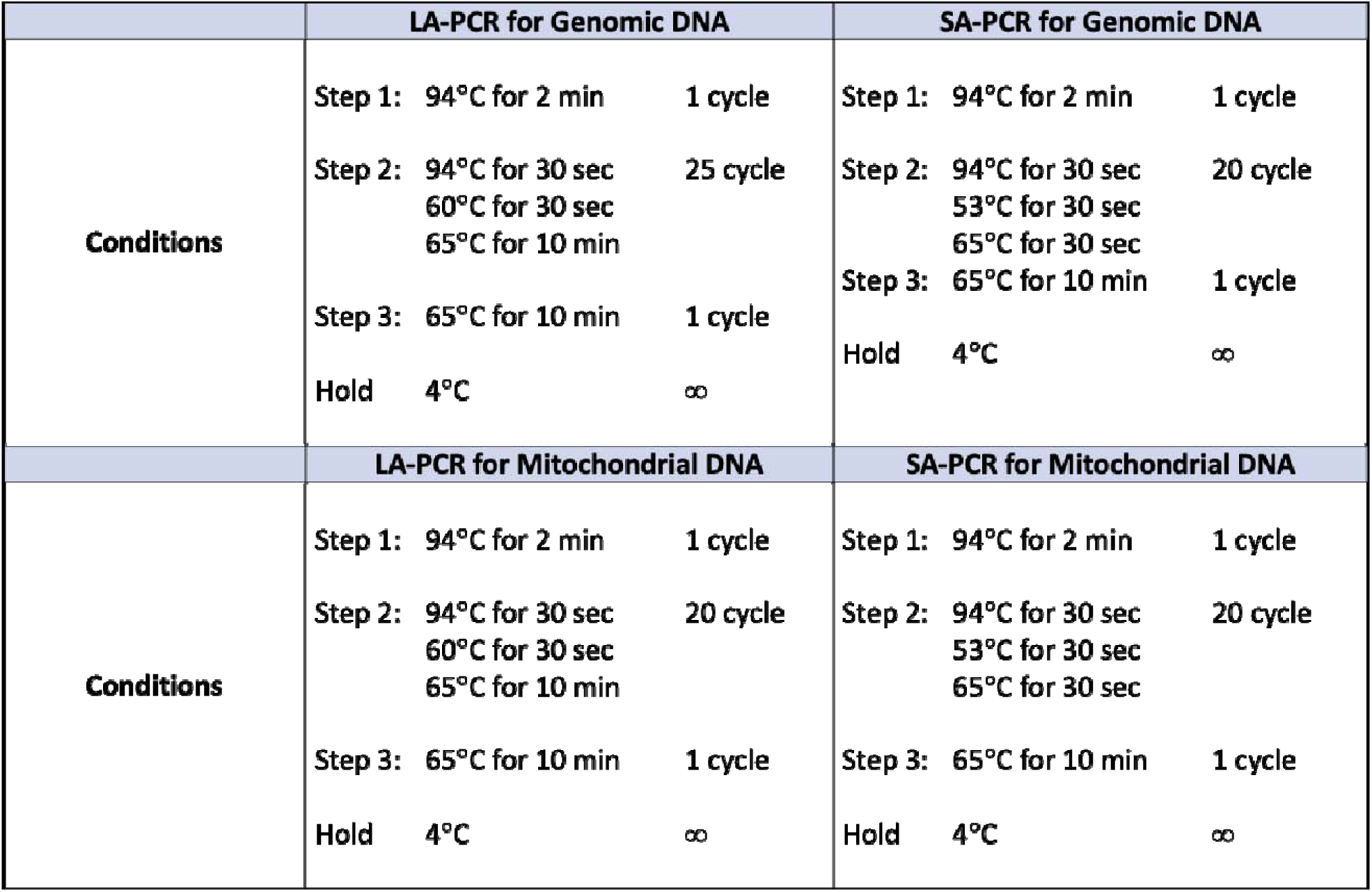
Primers and PCR program for LA-PCR.

## Acknowledgments

RNA core facility at HMRI for providing hTERT and GFP mRNA.

## Author contributions

D.F.C., A.M., and J.P.C. conceived the study and designed the experiments. A.M. and S.L. wrote the manuscript. T.K.C.N. and V.V.S. constructed libraries for RNA sequencing. V.V.S. and S.L. performed RNA sequencing analysis. K.C.P. made nanoparticles. D.F.C., A.M., S.L., E.M., A.T.L., J.C., performed experiments and collected data. J.P.C., K.W.B., A.J.S., O.D., and B.G. provided administrative support. All authors contribute and approve the final version of the manuscript.

---

**Supplementary Figure 1.**
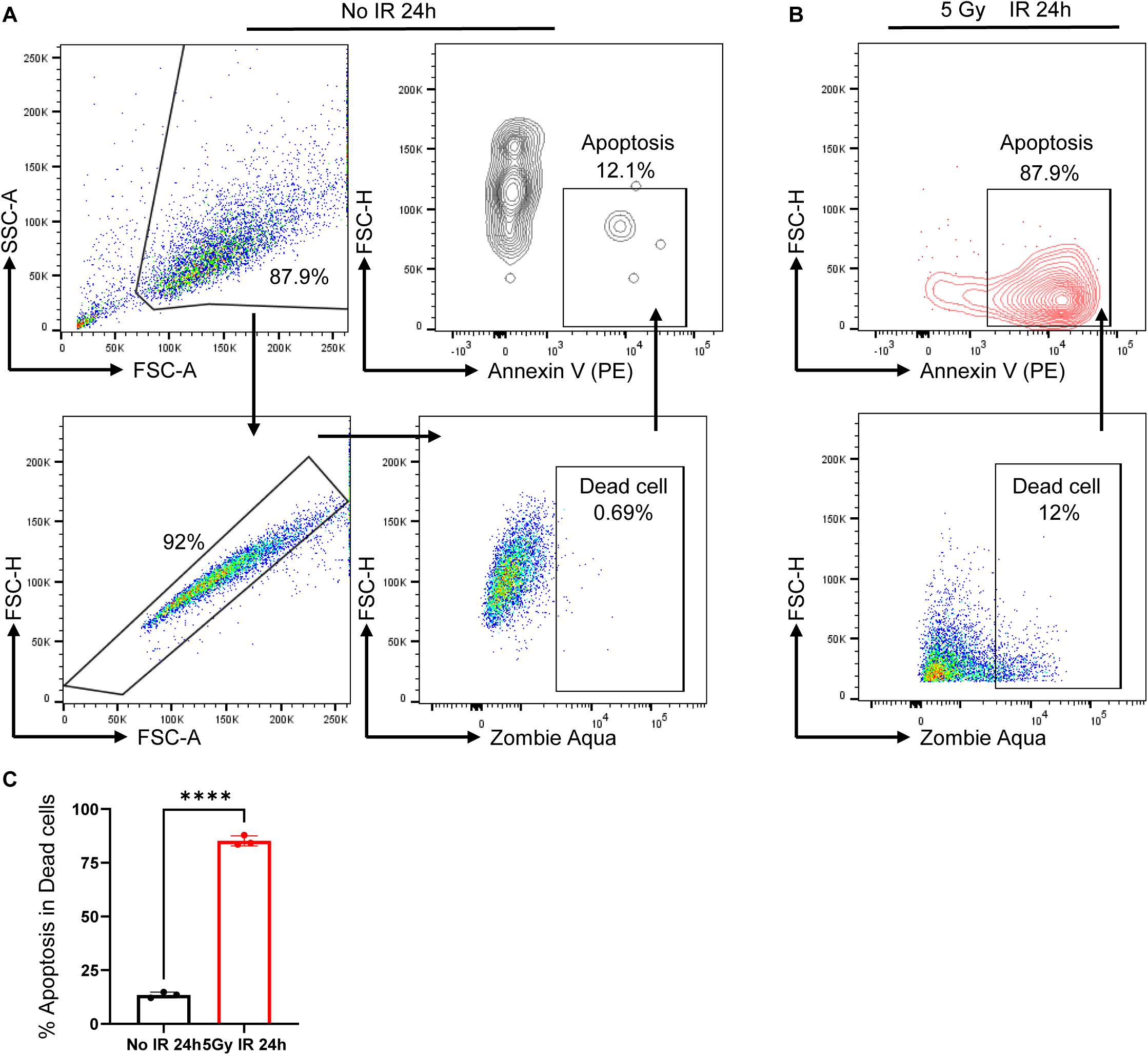
X-ray irradiation induces apoptosis in keratinocytes. **(A**) Gating strategy for flow cytometric analysis of cell apoptosis in keratinocytes. (**B**) Flow cytometry analysis reveals the percentage of apoptotic cells among dead keratinocytes 24 hours after exposure to 5 Gy irradiation. (**C**) Quantitative analysis of the percentage of apoptotic cells among dead keratinocytes, with or without 5 Gy irradiation, 24 hours post-treatment.

**Supplementary Figure 2.**
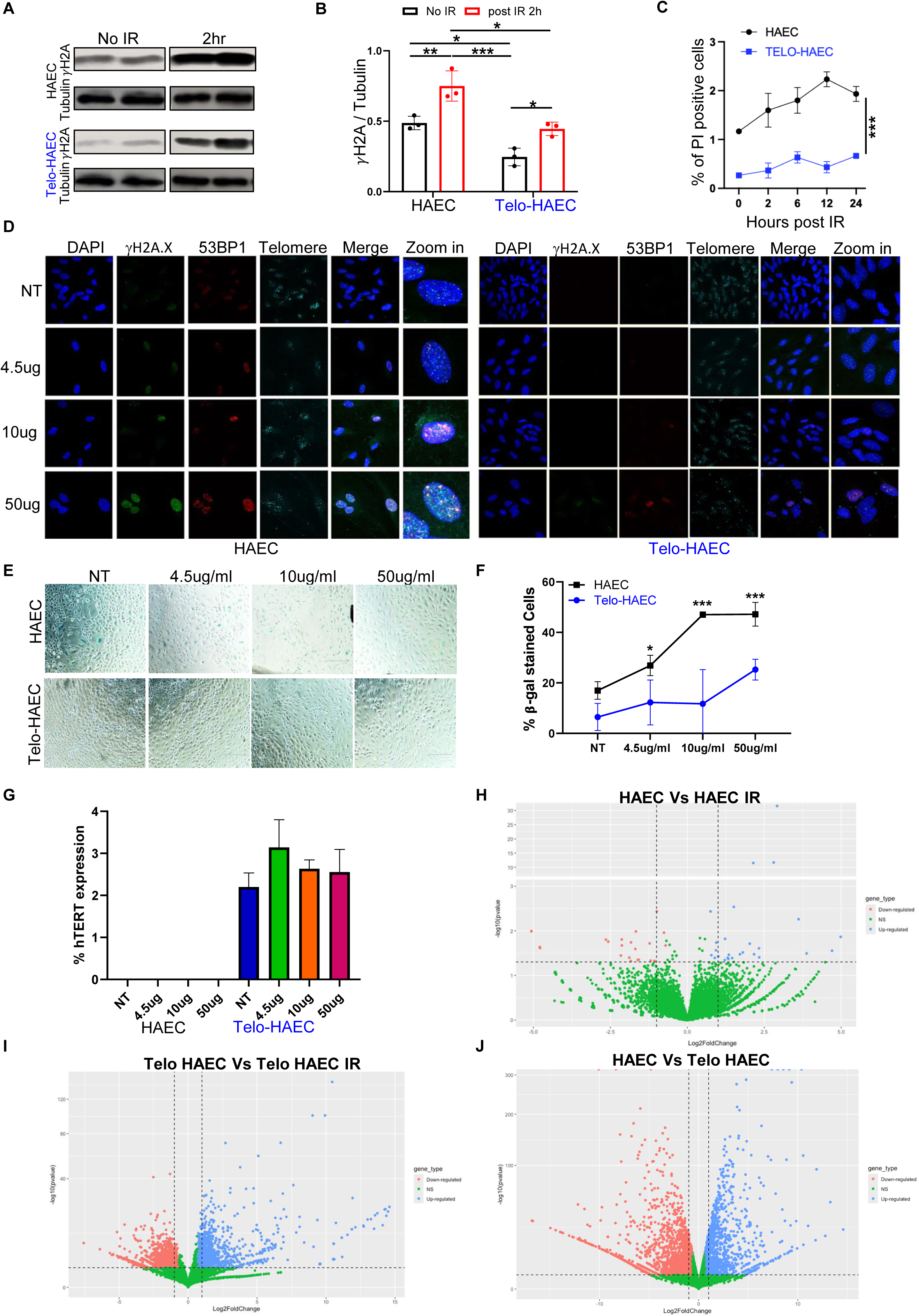
TERT overexpression in HAECs reduces DNA damage and protects cells from bleomycin-induced senescence. (**A-B**) Representative western blots and densitometric quantification of γH2AX protein levels post-radiation. (**C**) Quantification analysis of Propidium Iodide (PI)-positive cells, indicative of cell death, following radiation exposure. (**D**) Representative immunofluorescence staining showing a dose-dependent increase in γH2AX and 53BP1 foci in HAECs and Telo-HAECs treated with bleomycin. (**E, F**) Representative images and quantitative analysis of SA-β-gal-positive cells in HAECs treated with varying doses of bleomycin. (**G**) qPCR analysis of TERT expression in HAEC cells treated with different doses of bleomycin. (**H-J**) Volcano plots illustrate differentially expressed genes in (**H**) HAEC vs HAEC IR, (**I**) Telo-HAEC and Telo-HAEC IR, and (**J**) HEAC vs Telo HEAC.

**Supplementary Figure 3.**
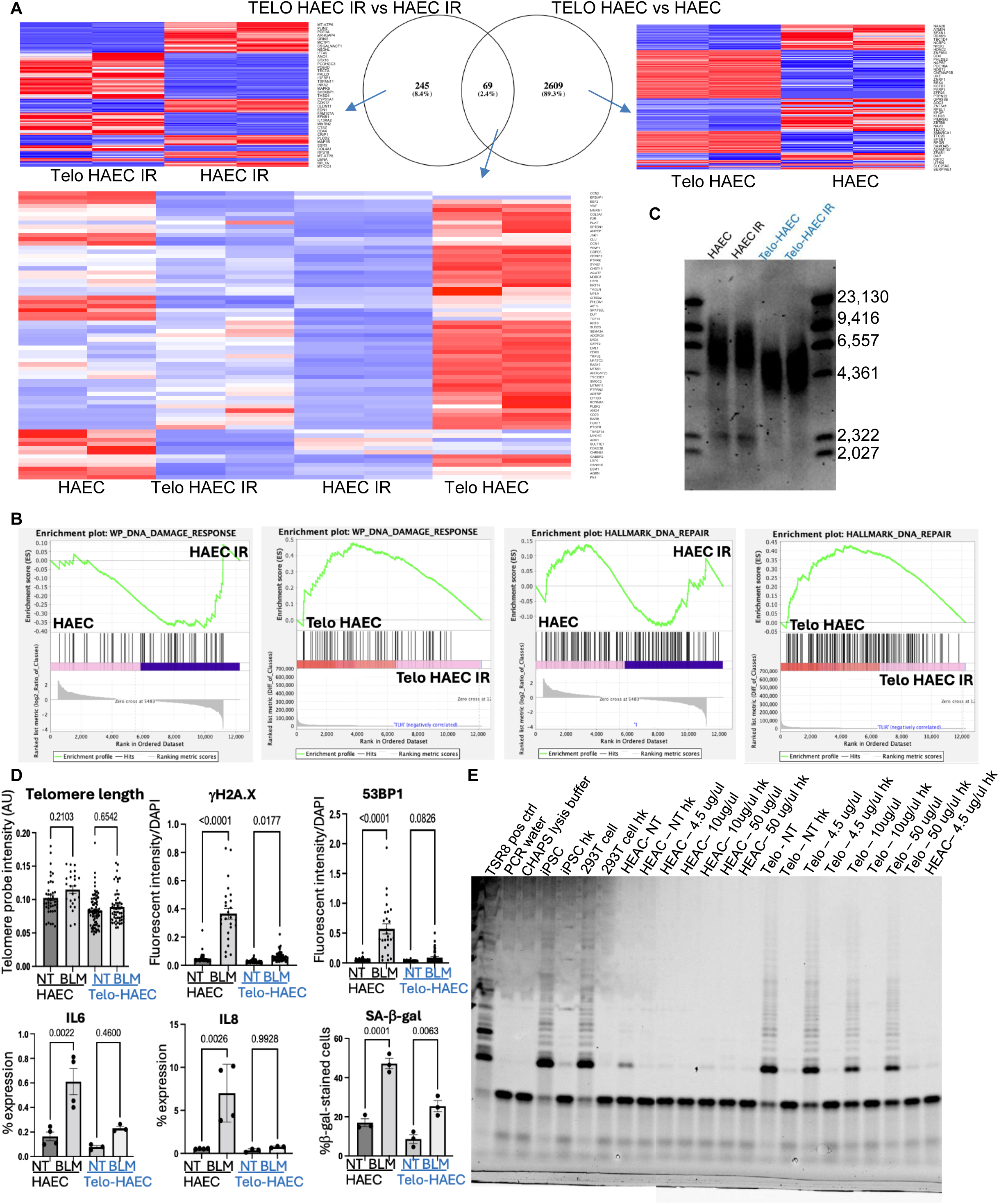
TERT overexpression in HAEC reduces DNA damage and increases cell survival. (**A**) Venn plot depicte pathways associated with differential expressed genes among HAEC-IR vs Telo-HAEC-IR and HAEC vs Telo-HAEC. Heatmap show gene differential expressed genes and common genes from Venn plot in HAEC-IR, Telo-HAEC-IR, HAEC, and Telo-HAEC groups. (**B**) Gene Set Enrichment Analysis (GSEA) highlights pathway enrichment for DNA damage response and DNA repair in comparisons between HAEC and HAEC IR, as well as Telo HAEC and Telo-HAEC IR. **(C**) Telomere length analysis by TRF (Terminal Restriction Fragment) assay in HAEC and Telo-HAEC 2h post-radiation. (**D**) qFISH analysis and quantification of γH2A.X, 53BP1, IL-6, IL-8, and SA-β-gal post-bleomycin treatment. (**E**) TRAP assay and qPCR analysis of TERT activity in addition and expression in HAECs and Telo-HAECs following bleomycin treatment.

**Supplementary Figure 4.**
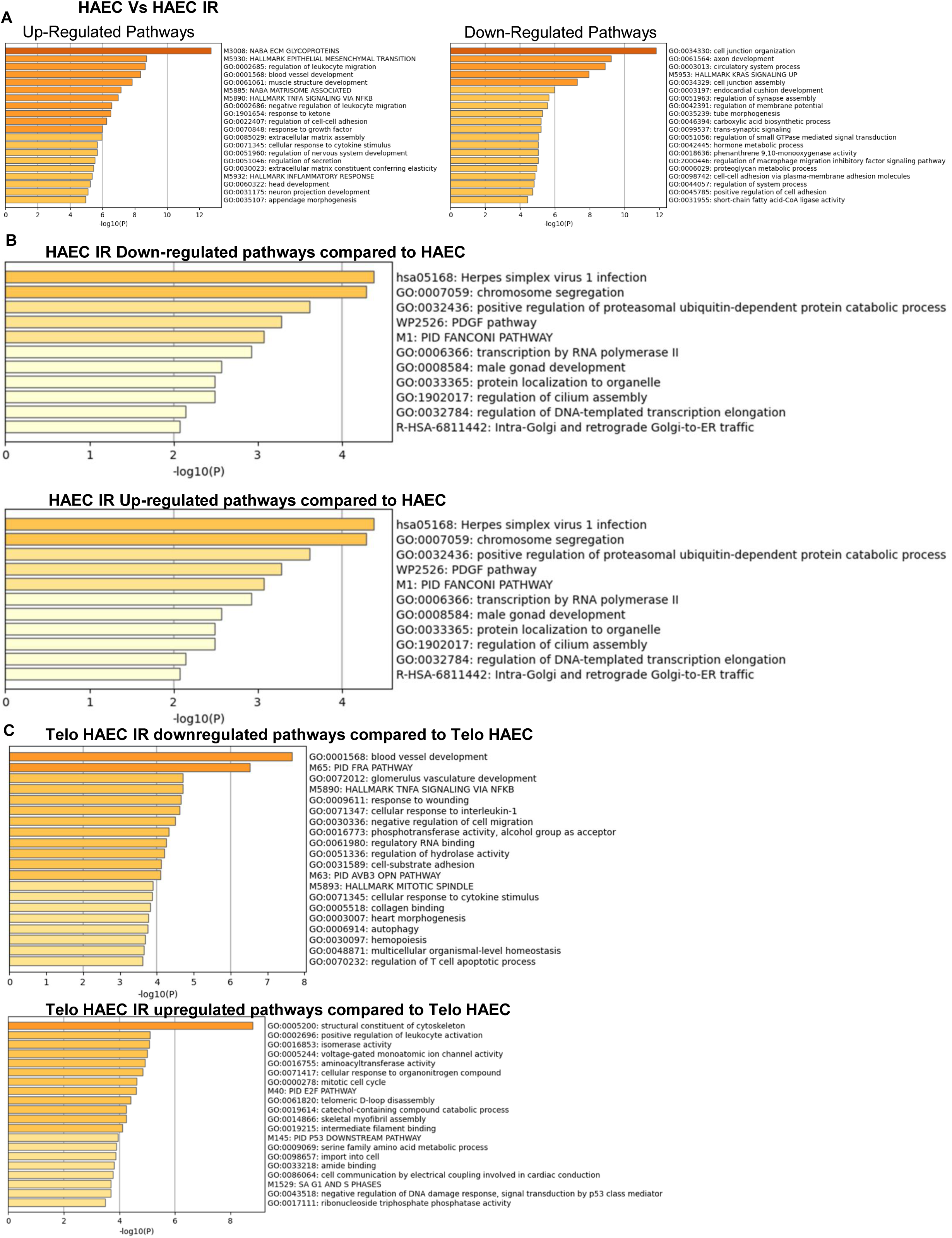
Pathway analysis was performed to compare different groups using differentially expressed genes identified from bulk RNA-seq data.

**Supplementary Figure 5.**
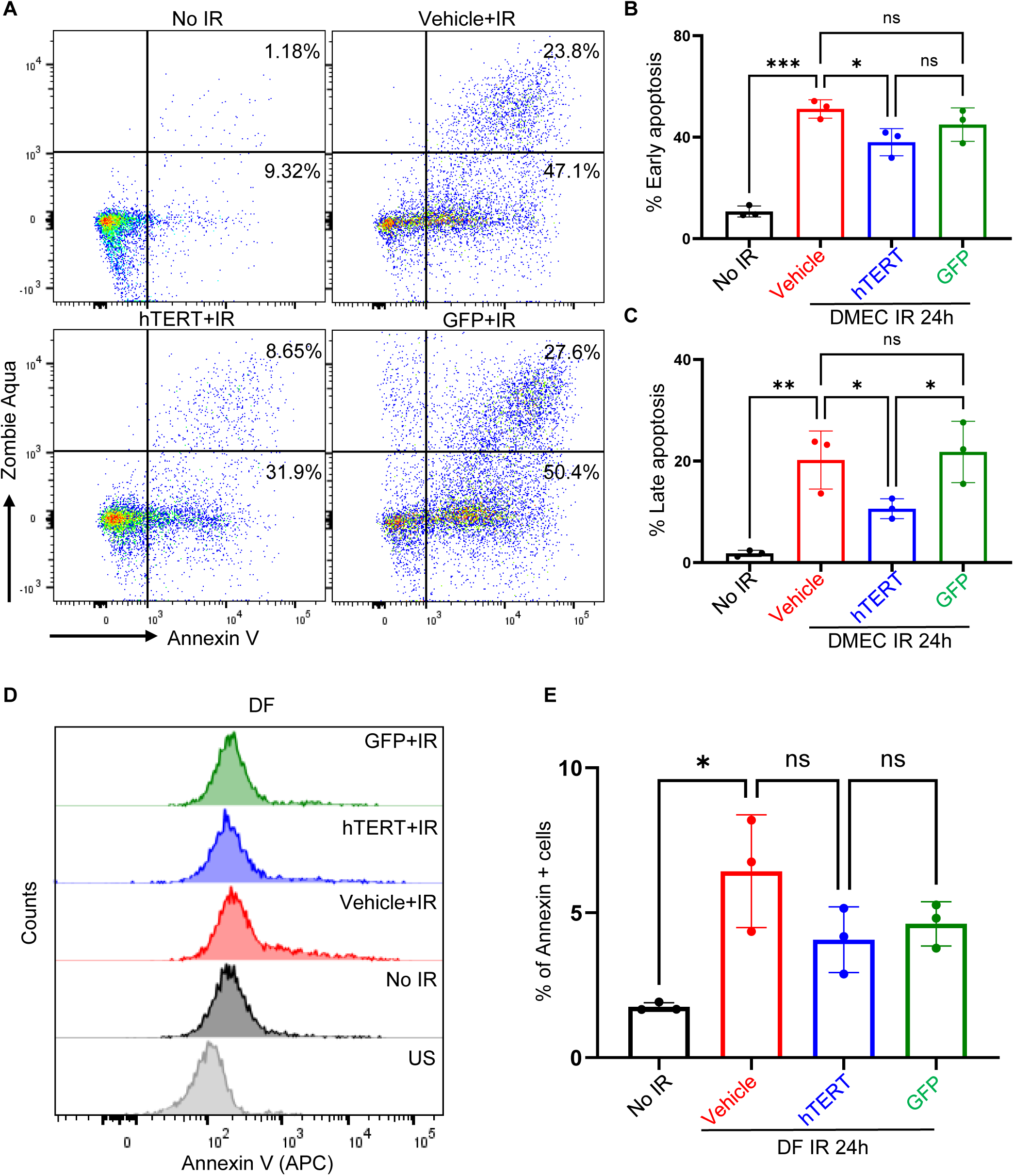
TERT mRNA treatment reduces radiation-induced apoptosis in primary skin cells. (**A-C**) Representative flow cytometry plots and quantitative analysis of early and late apoptosis in human dermal microvascular endothelial cells, assessed 24 hours after ionizing radiation (5 Gy) or no irradiation (No IR control). (**D, E**) Representative flow cytometry plots and quantitative analysis of Annexin V-positive cells in indicated groups of human fibroblasts. Data are shown as mean ± SD (n = 3). ns, *P* > 0.05; *, P < 0.05; **, *P* < 0.01; ***, *P* < 0.001. *P* values were calculated using two-way ANOVA. DMEC: Dermal Microvascular Endothelial Cells; DF: Dermal Fibroblasts.

